# Severe perinatal hydrops fetalis in genome edited pigs with a biallelic five base pair deletion of the Marfan syndrome gene, *FBN1*

**DOI:** 10.1101/2020.07.20.213108

**Authors:** Hiu-Gwen Tsang, Simon Lillico, Christopher Proudfoot, Mary E. B. McCulloch, Greg Markby, Violeta Trejo-Reveles, Brendan M. Corcoran, C. Bruce A. Whitelaw, Vicky E. MacRae, Kim M. Summers

## Abstract

This paper describes a genome editing project using CRISPR-Cas9. The objective was to create a large animal model of human Marfan syndrome by targeting the *FBN1* gene of the pig, *Sus scrofa*, using a single guide and non-homologous end joining which was expected to create short insertion or deletion mutations at the 5’ end of the gene. The editing successfully created a five base pair deletion in exon 2 of *FBN1*, which was homozygous in two animals. However, the phenotype of these piglets was unexpected, since they showed none of the signs consistent with Marfan syndrome but both suffered extreme hydrops fetalis with a large amount of fluid located under the skin and in the abdomen. One of the edited piglets was stillborn and the other was euthanised at birth on welfare grounds. It is likely that this result was due to unanticipated on- or off-target mutations, possibly in the *GLDN* gene 3 megabases away from *FBN1.* This result provides more evidence for unexpected outcomes of CRISPR- Cas9 gene editing and supports the proposal that all genome edited individuals should be subjected to strategies to track the CRISPR footprint, such as whole genome sequencing, before being used for further experimentation or in clinical applications.

## Introduction

Marfan syndrome (MFS; MIM 154700) is a genetic condition affecting connective tissue, most seriously the cardiovascular, ocular and skeletal systems (Aoyama et al., 1993; Judge and Dietz, 2005; Arslan-Kirchner et al., 2010). It is caused by mutations in the *FBN1* gene (MIM 134797) encoding the extracellular matrix fibrillar protein fibrillin-1. *FBN1* mutations are associated with a wide range of phenotypic effects affecting the connective tissue, including dilatation and dissection of the ascending aorta, ocular lens subluxation and dislocation and overgrowth of the long bones resulting in disproportionately long arms, legs and digits (Summers et al., 2006; Summers et al., 2012). Some mutations, primarily at the C-terminal end of the protein, result in lipodystrophy which may be associated with short stature, and a 140 amino acid peptide cleaved from the C-terminal acts as a glucogenic hormone (Romere et al., 2016; Davis et al., 2016). In humans, most pathogenic mutations are missense mutations (Collod-Beroud et al., 2003), often removing or introducing a cysteine in either the multiply repeated EGF domains or the highly conserved TGFβ-binding domains which are found only in three fibrillins and the four closely related latent transforming growth factor binding proteins (LTBPs) (Davis and Summers, 2012). Pathogenic deletions have also been found (Collod-Beroud et al., 2003; Li et al., 2017).

The major cause of mortality in MFS is dissection of the ascending aorta (Goyal et al., 2017). This is usually (but not always) preceded by dilatation of the aortic root, and lifespan of MFS patients has been increased by prophylactic surgery in which the root and the aortic valve are replaced (Milewicz et al., 2005). Further surgery may then be required as more distal parts of the aorta can dilate and dissect (Finkbohner et al., 1995). Surgery may also be possible for the subluxated lens and is essential for the detached retina which is associated with the long axial length of the globe (Nemet et al., 2006). With the extension of the lifespan other problems have been detected such as scoliosis-related morbidity, dissection of other arteries, long term ocular problems and other issues.

Other than management of the signs and symptoms by surgery, physiotherapy and lifestyle modifications, there are few treatment options for MFS (Ramirez et al., 2017). The aortic dilatation can be slowed using beta blocker drugs and the angiotensin pathway inhibitor losartan has shown limited promise (Takeda et al., 2016). The psychosocial aspects of diagnosis with a painful and life-threatening genetic condition which cannot be treated or prevented are severe, especially for young people who may have already lost a parent or grandparent to the disease (reviewed by Velvin et al., 2015). An animal model is needed to explore treatment options and to understand the implications of the condition as ageing progresses.

Mouse models of MFS have been important in understanding the pathophysiology of MFS (Pereira et al., 1997; Judge et al., 2004; Carta et al., 2006; Lima et al., 2010; Cook et al., 2012; Wilson et al., 2016), but have some limitations due to the small size of the mouse, the differences in structure between the mouse and human eye and the inconsistent genetic model, as many of the mouse phenotypes are recessive in contrast to the dominant human disease. A large animal model would overcome a number of these problems and allow more investigation of the phenotype with ageing. Natural bovine models have been reported for two cattle breeds, and bear strong resemblance to the human disease, with lens subluxation, aortic dilatation and skeletal abnormalities in heterozygous animals (Gigante et al., 1999; Singleton et al., 2005; Hirano et al., 2012). Pigs have been important in developing surgical techniques for cardiovascular problems, including heart transplants (Tsang et al., 2016), and would be an ideal model animal for further research into the MFS phenotype. The transcriptome of the pig across a wide range of organs and cell types has recently been published, to aid in the understanding of the pig model (Summers et al., 2019a). The pig genome sequence has recently been updated (Warr et al., 2020). In addition, we have looked at *FBN1* expression in the sheep, which reveals the high expression in cardiovascular structures, particularly the cardiac valves (Clark et al., 2017; Tsang et al., 2020).

*FBN1* edited pigs have been produced using the zinc finger nuclease approach, resulting in a single base deletion in exon 10 of *FBN1* (Umeyama et al., 2016). Heterozygous pigs had many manifestations of human Marfan syndrome and homozygous progeny showed neonatal lethality. In this project we aimed to create a pig model of Marfan syndrome using CRISPR-Cas9 genome editing and we anticipated a similar phenotype to that seen by Umeyama et al. (2016) for heterozygous and homozygous edited pigs.

## Methods

### Animal ethics

All animal work was approved by The Roslin Institute’s and the University of Edinburgh’s Protocols and Ethics Committees. The experiments were carried out under the authority of U.K. Home Office Project Licenses PPL60/4518 and PPL60/4482 under the regulations of the Animal (Scientific Procedures) Act 1986.

### Design and in vitro assessment of efficiency of guide RNAs

The online CRISPR Design tool (http://crispr.mit.edu/) was used for designing sgRNA. Potential sgRNA sequences were obtained for porcine *FBN1* (NCBI accession number NM_001001771.1; Ensembl transcript ID: ENSSSCT00000005143.4). Exon 2 was targeted. CRISPR guide sequences (approximately 20 nucleotides) were given by the tool, including information on likelihood of off-target cleavage. The plasmid vector used was pSL66, a derivative of px458 (Burkard et al., 2017). Cutting efficiency of each guide was assessed using porcine kidney PK15 cells, as described previously (Burkard et al., 2017). Based on *in vitro* cutting efficiencies, the guide chosen was FBN Guide 1-1 (5’-caccGGATGGAAAACCTTACC-3’), targeting the sequence GGATGGAAAACCTTACCTGG, where TGG is the protospacer adjacent motif (PAM) site (**Figure 1**).

**Figure 1.**
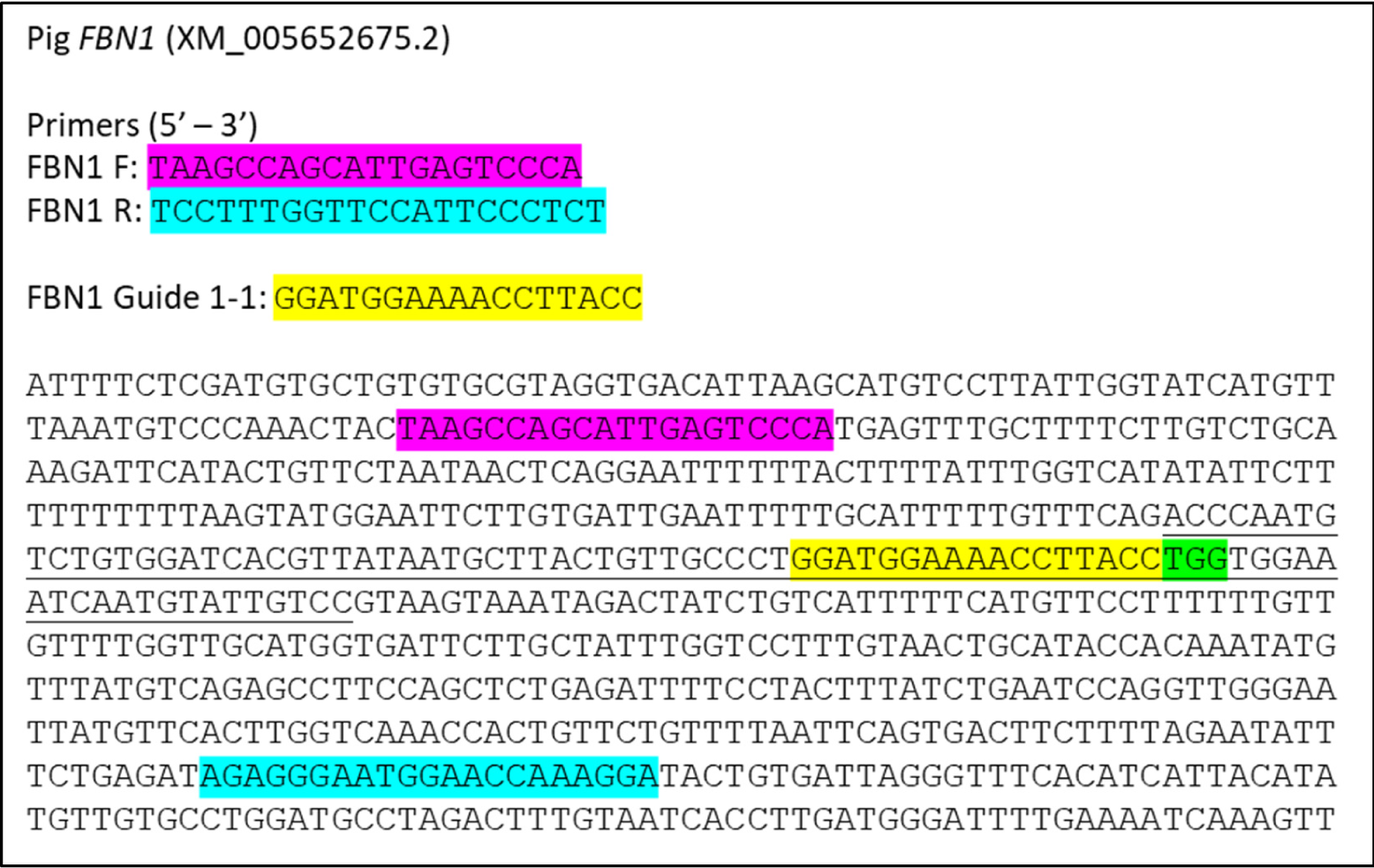
Porcine FBN1 sequence showing Exon 2 (underlined), the guide sequence (yellow), the PAM motif (green) and the primers used for genotyping (purple and blue).

### Generation of guide RNA and zygote injection

RNAs for zygote injection were generated by *in vitro* transcription and assessed as described (Burkard et al., 2017). The forward primer was 5’-ttaatacgactcactatagggGGATGGAAAACCTTACC-3’, which contains the T7 promoter (lower case) followed by the target sequence (upper case) and the reverse primer was oSL6 5’-aaaagcaccgactcggtgcc-3’ (Burkard et al., 2017). Zygote creation and injection and embryo transfer into host mother sows were performed as described previously (Lillico et al., 2013; Burkard et al., 2017). Following microinjection, pregnancy was confirmed at 4 weeks using ultrasound (performed by Dryden Farm Services, Roslin, UK).

### Genotyping of piglets

Ear and tail biopsies were taken from each newborn piglet for genotyping. DNA was extracted using the DNeasy Blood and Tissue kit for Cultured Cells (Qiagen, Manchester, UK) following the manufacturer’s protocol for the purification of total DNA from animal tissues. Samples were digested with proteinase K for 5 h at 56°C. DNA was subjected to PCR using Quickload Taq 2X mastermix (New England Biolabs, Hitchin, UK). The reaction mixture contained 25 μL Quickload mastermix, 1 μL forward primer (10 μM), 1 μL reverse primer (10 μM), 5 μL DNA (<1000 ng) and 18 μL nuclease-free water for a total volume of 50 μL. The forward primer was 5’ TAAGCCAGCATTGAGTCCCA 3’ and the reverse primer was 5’ TCCTTTGGTTCCATTCCCTCT 3’ (**Figure 1**). The PCR settings were as followed: 95°C for 30 s, 30 cycles of 95°C for 20 s, 60°C for 1 min and 68°C for 30 s, and a further 68°C for 5 min. PCR products were purified using the Chargeswitch PCR Clean-up kit (Thermo Fisher Scientific, Waltham, MA USA), before quantifying with a NanoDrop spectrophotometer (Thermo Fisher Scientific) and sequencing. Samples were sent for chain termination sequencing by Edinburgh Genomics (University of Edinburgh, UK). Sequencing data were analysed using SeqMan Pro and SeqBuilder (Lasergene 13, DNASTAR Inc, Madison, WI, USA).

### Post mortem examination

Post mortem examination of genome edited pigs was performed by coauthors McCulloch and Summers, within 48 hours of birth. The eyes were dissected and stored in Davidson’s Fixative. Computerised tomography (CT) scanning using a Somatom Definition AS (Siemens Healthcare Ltd, Surrey, UK) was performed by the Large Animal Imaging Service at the Large Animal Unit, Dryden Farm, Edinburgh, UK. Resulting images were viewed in the OsiriX programme (Pixmeo SARL, Bernex, Switzerland).

## Results

### sgRNA guide designs

sgRNAs were designed for porcine *FBN1* using the CRISPR Design tool (http://crispr.mit.edu/). Within the output, guide sequences with PAM sites, required for targeting by Cas9 endonuclease, were given. Quality scores (inverse likelihood of off-target binding) were provided, where higher scores were preferable. These guide sequences were modified to ensure that they possessed a 5’ guanine (G) nucleotide. Two guides were tested in PK15 cells. The guide chosen had the highest quality score (77), no predicted off target sites in genes and a cutting efficiency of 40% in the T7 endonuclease assay.

### G*enome edited piglets*

The CRISPR-Cas9 construct was then introduced by microinjection into pig zygotes harvested from donor sows. 2 – 5 pL of the injectate containing the guide RNA and Cas9 mRNA were delivered to the cytoplasm. The zygotes were then injected into the oviducts of a nulliparous recipient sow. At 4 weeks the sow was scanned and pregnancy with at least four foetuses was confirmed. At full term (16 weeks) two healthy piglets were delivered (**Figure 2A**) followed by a live born piglet with major abnormalities and a still born piglet with the same abnormal phenotype (**Figure 2B**). The still born piglet was felt to be recently deceased as there were no signs of necrosis or mummification and the phenotype was identical to its live born affected littermate. These two piglets showed excessive oedema, particularly affecting the abdomen, neck and limbs. The skin appeared shiny and thin. The liveborn piglet took some breaths and attempted to walk, but was unable to support its weight on the hindlimbs. It was euthanised immediately after birth on welfare grounds.

**Figure 2.**
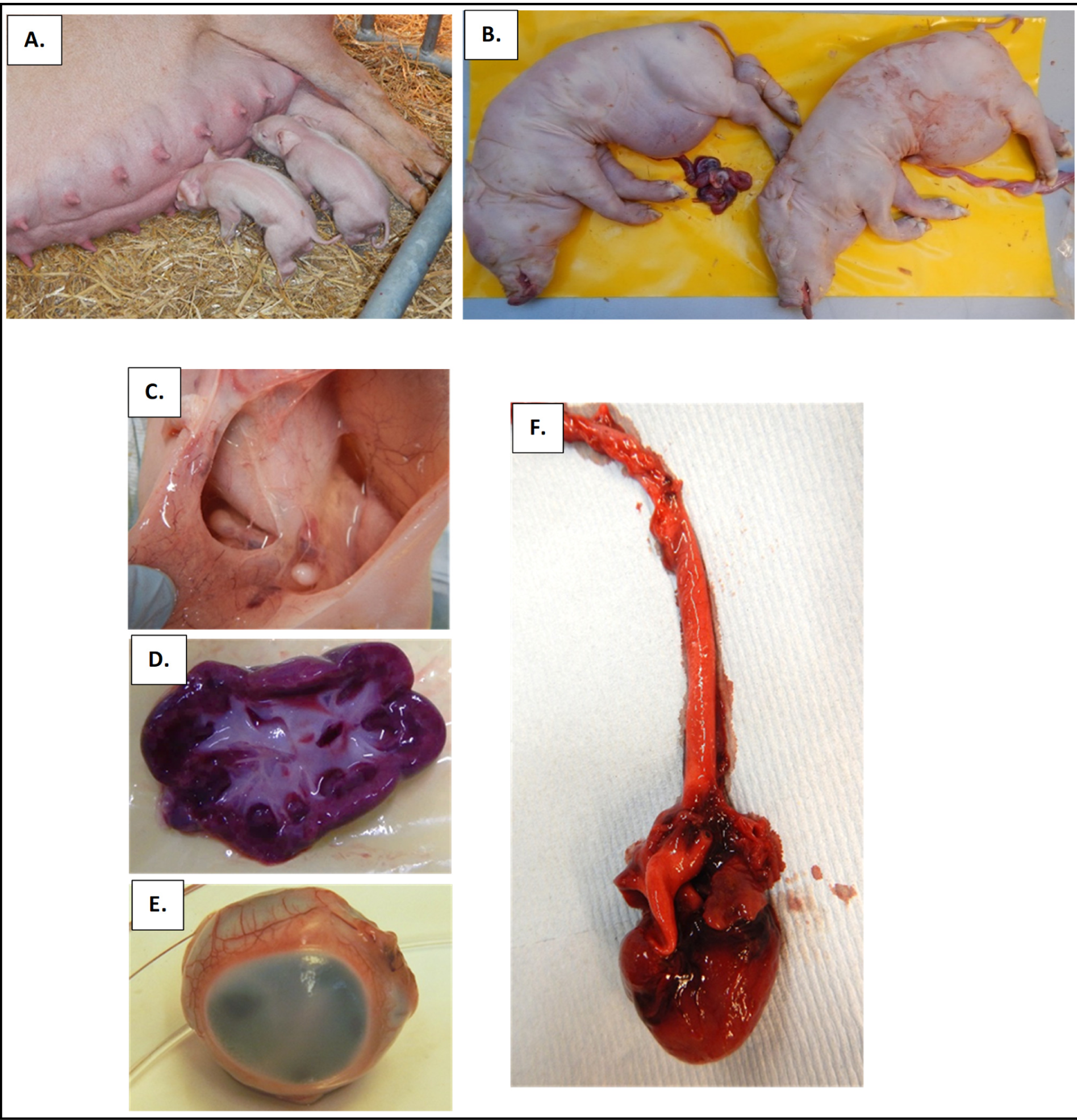
Genome editing of piglets. **A.** Healthy liveborn piglets suckling after birth. **B.** Piglets with homozygous 5 bp deletion in *FBN1*. Left, liveborn female piglet, euthanised immediately after birth; right stillborn male piglet. **C.** Fluid beneath and within the skin. **D.** Large, irregularly lobulated kidney. **E.** Eye, showing opaque cornea. **F.** Heart and aorta.

The day after birth the piglet cadavers were subjected to CT scanning. There were no obvious abnormalities. The ocular lenses appeared to be in place, there was no obvious scoliosis and the limbs appeared to be in proportion. Postmortem examination (within 48 hours after birth) confirmed the presence of clear colourless fluid in the skin, abdomen, legs and feet (**Figure 2C**) of both anomalous piglets. The skin had reduced hair, probably related to the oedema. The spine was not obviously scoliotic and the limbs were not elongated. There was a normal oral cavity with teeth erupted as normal and there was no meconium. The linea alba was intact in both piglets indicating that there was no body wall defect and there were no adhesions in the abdomen. Joints were relatively normal although one hip joint in each animal had restricted movement. The kidneys appeared large and irregularly lobulated (**Figure 2D**). The diaphragm was thin in the centre (around the oesophageal hiatus) but muscular at the edges. The corneas of all four eyes were opaque (**Figure 2E**). The heart appeared normal (**Figure 2F**) and there was no blood in the chest or abdominal cavity indicating that there had not been an aortic rupture. The cardiac valves showed some indication of myxomatous degeneration at the tips of the leaflets. The brain was friable but normal in appearance.

### Genotyping of piglets

DNA extracted from ear tissue of all piglets was amplified for the targeted region of the *FBN1* gene (**Figure 1**) and the amplicons sent for chain termination sequencing. The two healthy piglets were determined to be homozygous for the normal allele. The still born and euthanised piglets were homozygous for an identical 5 base pair deletion of the *FBN1* gene (**Figure 3**).

**Figure 3.**
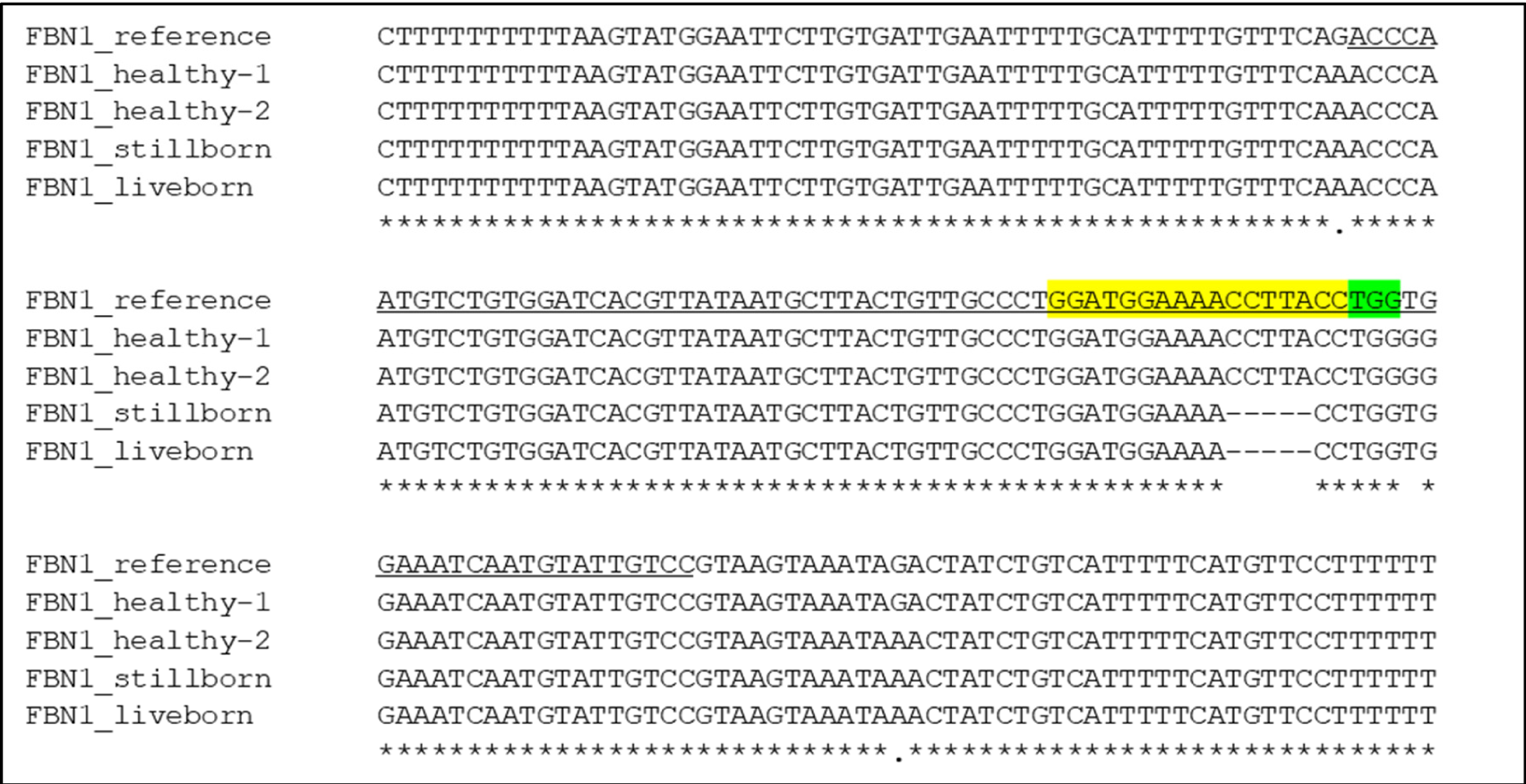
Sequence of Exon 2 and adjacent intronic sequence in reference, healthy and affected piglets. Guide sequence highlighted in yellow, PAM sequence in green and exon 2 underlined in the reference sequence. The five-base pair deletion is within the guide sequence, in both affected piglets.

## Discussion

Hydrops fetalis is a condition in which fluid builds up in the organs and tissues of a foetus. In humans in the past it was most commonly seen as a consequence of Rhesus blood group (Rh) incompatibility between foetus and mother (haemolytic disease of the newborn), but more recently detection and management of Rh negative mothers has minimised this and hydrops fetalis is now commonly seen due to anaemia, infection at birth, heart, lung or liver defects. It occurs in about 1 per 1000 human births. 50% of human infants affected with hydrops fetalis do not survive (Nassr et al., 2018; Yeom et al., 2015). The Online Mendelian Inheritance in Man data base lists a number of genetic conditions in which hydrops fetalis may be a feature (**Table 1**). Notably, in many of these conditions there is a skeletal abnormality (Abboy et al., 2008). In South East Asia hydrops is commonly associated with alpha thalassemia and the production of the abnormal haemoglobin Hb Bart (haemoglobin gamma tetramers, which have high oxygen affinity and are incapable of releasing oxygen in the periphery). The incidence is as high as 1 in 200 in some regions (King and Higgs, 2018). Hydrops fetalis has also been associated with Noonan syndrome (mutations of *RIT1* or *RAF1* gene) (Yaoita et al., 2016; Australian Genomics Health Alliance Acute Care Flagship, 2020), with a glycosylation disorder (*PMM2* gene; Panigrahy et al., 2016) and with biallelic mutations of *LARS2* and *GLDN* (Australian Genomics Health Alliance Acute Care Flagship, 2020). Hydrops fetalis has also been reported in five animal species of a number of breeds (**Table 2**). No genes have been associated with the animal cases, although several affected lambs were related through the maternal grandfather and/or father, suggesting a possible genetic aetiology.

**Table 1.** Conditions with hydrops fetalis recorded in Online Mendelian Inheritance in Man (OMIM) (https://omim.org). ^a^ Patient reported by Yaoita et al. (2016). ^b^Patients reported by Australian Genomics Health Alliance Acute Care Flagship (2020).

| Condition | OMIM # of condition | Gene | OMIM # of gene | Location |
| --- | --- | --- | --- | --- |
| Gaucher disease, perinatal lethal | 608013 | *GBA* | 606463 | 1q22 |
| Noonan syndrome 8 | 615355 | *RIT1*<sup>a</sup> | 609591 | 1q22 |
| Greenberg dysplasia | 215140 | *LBR* | 600024 | 1q42.12 |
| Lymphatic malformation 8 | 618773 | *CALCRL* | 114190 | 2q32.1 |
| Hydrops, lactic acidosis, and sideroblastic anaemia | 617021 | *LARS2* | 604544 | 3p21.31 |
| Noonan syndrome 5 | 611553 | *RAF1*<sup>b</sup> | 164760 | 3p25.2 |
| Mucopolysaccharidosis Type VII | 253220 | *GUSB* | 611499 | 7q11.21 |
| Lymphatic malformation 7 | 617300 | *EPHB4* | 600011 | 7q22.1 |
| Lethal congenital contracture syndrome 11 | 617194 | *GLDN*<sup>b</sup> | 608603 | 15q21.2 |
| Congenital disorder of glycosylation Type 1a | 212065 | *PMM2* | 601785 | 16p13.2 |
| Hydrops fetalis, non-immune; Haemoglobin H disease | 236750; 613978 | *HBA1*<br>*HBA2* | 142800<br>141850 | 16p13.3 |
| Congenital pulmonary lymphangiectasia | 265300 |  |  |  |
| Nonimmune hydrops fetalis with gracile bones and dysmorphism | 613124 |  |  |  |

**Table 2.**
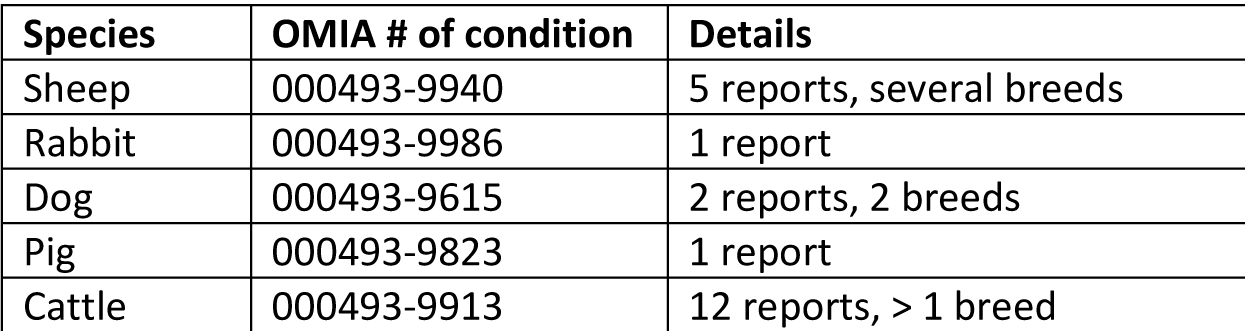
Conditions with hydrops fetalis recorded in Online Mendelian Inheritance in Animals (OMIA) (https://omia.org).

In human infants hydrops fetalis is associated with enlargement of the heart, liver and spleen, pale colouring and difficulty breathing. A report of hydrops fetalis in lambs noted severe generalised subcutaneous fluid accumulation and marked effusions in thoracic and abdominal cavities (Alleaume et al., 2012). This is similar to the two piglets produced here. It might be expected that animals homozygous for a truncating *FBN1* mutation would have cardiovascular abnormalities, and cardiac anomalies are commonly the cause of human hydrops fetalis (Yuan, 2017; Nassr et al., 2018). We did not notice any obvious pathology of the heart or aorta and there was no evidence of a dissection or rupture (no blood in the chest cavity). However, the presence of hydrops could be an indication of a cardiovascular abnormality. Post-mortem examination did not note gross abnormalities of any organs, although the kidneys were felt to be large and there were some structural abnormalities of the urethras and diaphragm.

The most striking pathology in the gene-edited piglets was the dense opacity of the cornea (**Figure 2E**), which has not been described in MFS. This opacity was more pronounced than the corneal clouding often seen in eyes after death. The lens appeared to be in place by both histological examination and CT scan. We did not observe obvious anomalies in limb length, spine or body size compared with the wild type litter mates, and CT scan did not reveal gross abnormalities consistent with MFS.

The previous pigs with a homozygous *FBN1* mutation (Umeyama et al., 2016) and the rare humans who have biallelic or homozygous pathogenic mutations did not show a phenotype of hydrops fetalis. The human patients showed a wide range in severity of Marfanoid features (Chemke et al., 1984; de Vries et al., 2007; Hogue et al., 2013; Arnaud et al., 2017). mRNA transcribed from the 5 bp deletion in these piglets would likely be susceptible to nonsense mediated decay (Baker and Parker, 2004) and if translated would result in a severely truncated protein, so that the piglets would have no functional fibrillin-1 protein. This may predispose to the hydrops phenotype since the human cases tend to be missense mutations (Arnaud et al., 2017). However, it is surprising that we saw no signs of MFS. Perhaps the lack of fibrillin-1 results in a less severe phenotype than the presence of a dominant negative abnormal protein. Alternatively, the manifestations of MFS, such as lens subluxation and aortic dilatation, are not always present at birth in heterozygous individuals and may have appeared in these piglets as they aged.

Hydrops was recently seen in an infant with biallelic mutations in the gliomedin gene, *GLDN*, located 3 megabases downstream of *FBN1* (Australian Genomics Health Alliance Acute Care Flagship, 2020; see **Table 1**). There have been a number of reports of DNA changes in the vicinity of a target sequence associated with the use of CRISPR-Cas9 gene editing (Summers et al., 2019; Alanis-Lobato et al., 2020; Liang et al., 2020; Zuccaro et al., 2020). It is possible that *GLDN* was mutated by the CRISPR-Cas9 process and is responsible for the hydrops fetalis phenotype in our piglets.

## Conclusion

We present a case of an unanticipated phenotype associated with CRISPR-Cas9 gene editing. We have previously reported the reinsertion of a deleted fragment close to the CRISPR target site in a mouse line (Summers et al., 2019b) and others have presented similar outcomes in edited human embryos, including loss of whole chromosomes (Alanis-Lobato et al., 2020; Liang et al., 2020; Zuccaro et al., 2020). It is not clear whether the phenotype in these piglets was due to an on-target event similar to these or an off-target event (although this was predicted to have a low probability by the guide design software), perhaps affecting the nearby *GLDN* gene, as we did not subject the DNA to whole genome sequencing. A number of strategies have been developed to track the CRISPR footprint in the edited genome (Lin and Luo, 2019). It would seem wise to use these strategies to examine the whole genome of edited individuals to confirm that the only genetic change is that predicted by the guides (Smith et al., 2014).

## Acknowledgements

The study was supported by Roslin Institute Strategic Program grant ‘Blueprints for Healthy Animals’ (BB/P013732/1) from the Biotechnology and Biological Sciences Research Council, UK. Mater Research Institute-UQ is grateful for support from the Mater Foundation, Brisbane. The Translational Research Institute receives core support from the Australian Government. HGT was supported by studentship funding via the East of Scotland BioScience Doctoral Training Partnership BB/J01446X/1 (EASTBIO DTP). MEBM received funding from the Royal (Dick) School of Veterinary Studies, University of Edinburgh. GM was funded by Dogs Trust, UK. VTR was supported by a CONACYT (Mexico) international studentship. The funders had no role in study design, data collection and analysis, decision to publish, or preparation of the manuscript.

